# A streamlined spectral cytometry method for FAD and NADH autofluorescence analysis in immunometabolic studies

**DOI:** 10.64898/2026.06.03.729953

**Authors:** Emmanouil Stylianakis, Nadine Hövelmeyer

## Abstract

We present a streamlined protocol that enables the characterization of the metabolic state of immune cell populations through their distinct NADH/FAD autofluorescence fingerprints using a FACSymphony A5 spectral cytometer. We demonstrate the utility of this approach by profiling the metabolic status of diverse splenic B-cell subsets and assessing metabolic changes associated with their activation state.

## Introduction

Immunometabolism has emerged as a rapidly evolving field that investigates how cellular metabolic programs regulate immune cell function across diverse contexts, including autoimmunity, infection, aging, and cancer. Immune cells rely primarily on two bioenergetic pathways, cytosolic glycolysis and mitochondrial oxidative phosphorylation (OXPHOS), to meet energetic and biosynthetic demands. Upon activation or in response to environmental challenges, metabolic pathways are dynamically reprogrammed to support proliferation, differentiation, survival, redox balance, effector functions, biosynthetic processes, and even cell fate decisions ^1,2^.

Established approaches for assessing cellular metabolism include systems-level techniques such as RNA sequencing and mass spectrometry-based metabolomics, as well as platforms such as the Seahorse XF analyser, which measure the oxygen consumption rate (OCR) and extracellular acidification rate (ECAR) as proxies for mitochondrial respiration and glycolysis, respectively. While these technologies provide valuable functional insights, they are limited by high cost, complex workflows, relatively high cell number requirements, and low scalability^3^. In particular, Seahorse-based OCR measurements do not account for mitochondrial abundance, making it difficult to distinguish whether increased respiration reflects enhanced activity per mitochondrion or increased mitochondrial number with similar electron transport chain activity. To address these limitations, various flow-cytometry-based approaches can be utilized. Voltage-dependent accumulating dyes, like TMRM, constitute an easy way to stain for active mitochondria, commonly in combination with dyes that stain mitochondrial mass (e.g., Mitotracker dyes). However, TMRM signals do not always align with Seahorse XF-derived oxygen consumption rate, since TMRM signals may be influenced by additional factors, such as metabolic stress, proton leak, reverse electron transport or oxidative stress^3–5^. SCENITH and Met-Flow comprise additional tools that enable metabolic analysis at the single-cell level within various populations ex and in vivo^6–8^. However, these methods primarily rely on protein abundance as a surrogate for metabolic activity, which may not accurately reflect functional metabolic states because protein abundance is influenced by numerous non-metabolic processes that regulate protein synthesis and degradation, such as ribosomal efficiency, proteasome activity or endoplasmic stress, which determine protein turnover, especially under chronic inflammatory or autoimmune settings.

An alternative robust approach is the measurement of endogenous autofluorescence of metabolic factors, which provides a direct, label-free readout of cellular metabolic activity. Nicotinamide adenine dinucleotide (NADH) and flavin adenine dinucleotide (FAD) are central electron carriers in the tricarboxylic acid (TCA) cycle and mitochondrial electron transport chain, transferring electrons to complexes I and II, respectively. Their redox states are therefore tightly coupled to mitochondrial function. Importantly, both NADH (reduced form of NAD) and FAD (oxidized form of FADH_2_) exhibit intrinsic autofluorescence, enabling real-time assessment of cellular metabolism without exogenous labelling ^9–11^. A low ORR, driven by high NADH fluorescence, reflects a reduced metabolic state typically associated with glycolytic activity, whereas a high ORR, driven by high FAD fluorescence, reflects an oxidized state associated with OXPHOS. Given the roles of NADH and FADH_2_ in mitochondrial electron transport, the ORR provides a functional proxy and normalized surrogate of mitochondrial respiratory activity, independent of cell size and mitochondrial abundance, which correlates with OCR measurements obtained using Seahorse platforms^11,13,14^.

A critical technical consideration identified in our analysis was the impact of excitation wavelength on accurate FAD detection. Previous studies have utilized the ultraviolet laser to excite FAD (Fig 1A, C). However, we illustrate that excitation of FAD in the UV range resulted in substantial spectral spill-over from NADH autofluorescence, leading to signal contamination, which makes it difficult to monitor FAD levels accurately (Fig. 1A, D, F). Under mitochondrial stress conditions, this UV-derived signal exhibited an unexpected trajectory, moving in parallel to NADH rather than the expected inverse relationship between NADH and FAD redox states^15^, indicating that it does not accurately represent FAD dynamics (Fig. 1A, B, D, F). Based on these findings, we strongly recommend excitation of FAD using blue or violet laser lines, rather than the ultraviolet one, to minimize spectral overlap and ensure reliable quantification, as these wavelengths do not excite NADH (Fig. 1G-H). In addition, by incorporating SSC-A, one may be able to distinguish metabolic activity driven by increased mitochondrial numbers or increased enhanced activity per mitochondrion ^16,17^.

**Figure 1:**
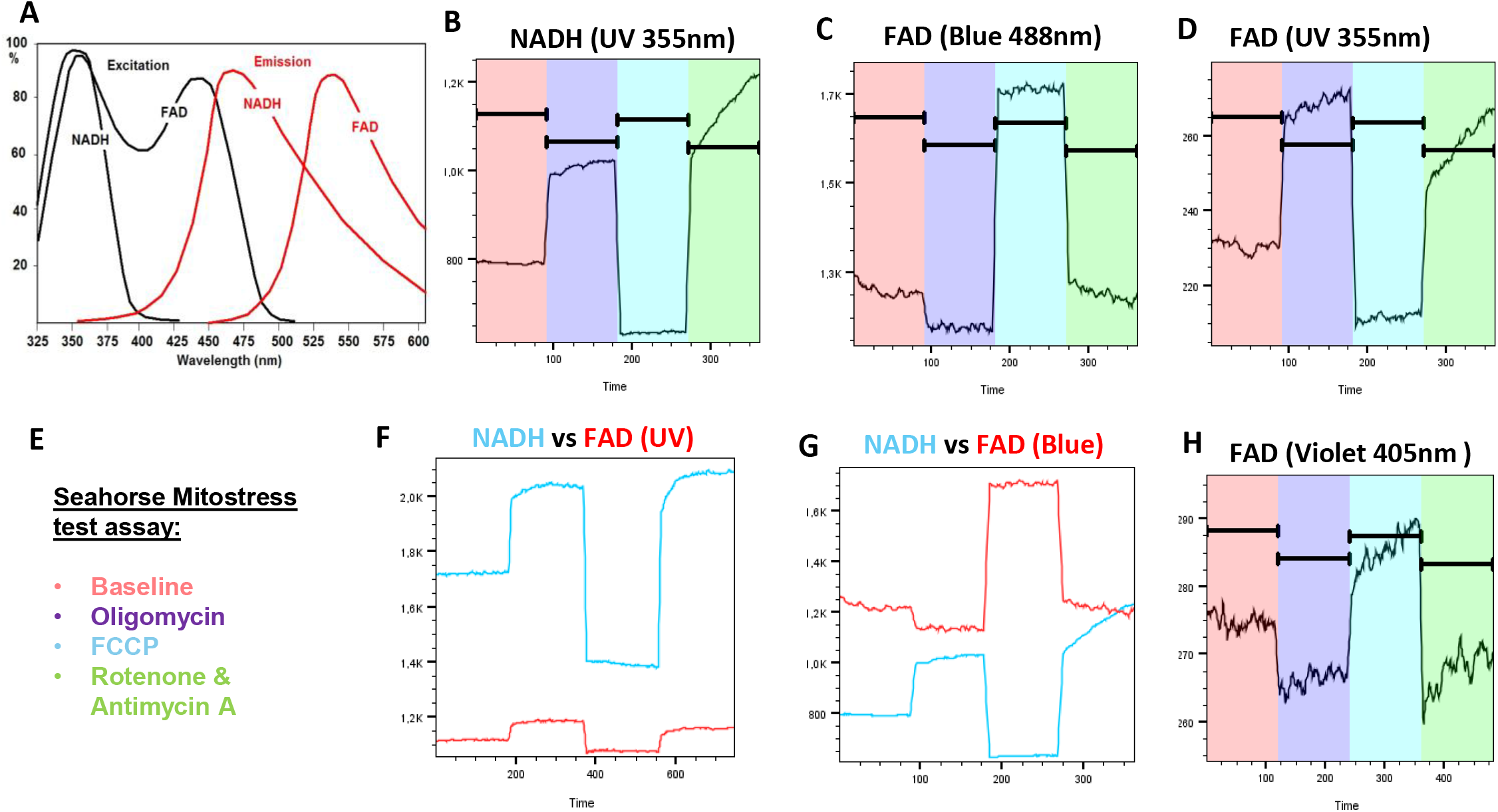
Real-time excitation and emission collection of NADH and FAD in the ultraviolet, violet, and blue region, at steady-state and following sequential treatment with oligomycin, FCCP, and rotenone/Antimycin A. (A) Excitation and emission spectra for NADH and FAD *(Adapted from Becker & Hickl: Metabolic imaging by FLIM)* illustrate a single excitation peak for NADH and two excitation peaks for FAD. (B) NADH was excited using a 355 nm ultraviolet (UV) laser, and emission was detected using a 446/67 nm band-pass emission filter and a 425 nm dichroic mirror. FAD excitation was assessed using (C) a 488 nm blue laser and emission was detected through a 537/32 nm band-pass emission filter using a 520 nm dichroic mirror, or (D) with a 355 nm UV laser, with emission being collected in the corresponding channels using appropriate bandpass filters. Application of the (E) mitostress assay allowed us to track NADH and FAD dynamics, (F) FAD detection using 355 nm excitation showed substantial spectral spill-over from NADH, limiting its specificity, as it changed in parallel with NADH. (G) In contrast, excitation with the 488 nm blue laser provided optimal separation. (H) FAD signal is also detectable upon excitation with a 405 nm violet laser, representing a second-best option, when, for instance, the green channel is occupied by GFP or another green fluorochrome.

To experimentally validate the responsiveness of NADH and FAD autofluorescence to mitochondrial perturbations, as measured using an ultraviolet (UV) and Blue laser, respectively, we adapted the Seahorse Mito Stress Test to monitor real-time NADH and FAD dynamics^5,11,15^. Pharmacological modulation of mitochondrial function induced predictable and opposing changes in NADH and FAD redox states. Inhibition of ATP synthase (complex V) with oligomycin reduced proton flux and increased mitochondrial membrane potential, resulting in the accumulation of reduced cofactors (NADH and FADH_2_). Subsequent treatment with the protonophore FCCP uncoupled electron transport from ATP synthesis, driving maximal electron transport activity and promoting oxidation of NADH to NAD and FADH_2_ to FAD. This resulted in decreased NADH fluorescence and increased oxidized FAD signal. Finally, inhibition of complexes I and III using rotenone and antimycin A halted electron transport, leading to NADH accumulation and depletion of oxidized FAD (Fig. 1B, C, G).

Considering that the optical redox ratio is a normalized parameter, it does not distinguish whether elevated metabolic activity arises from increased mitochondrial/electron transport chain activity or simply from greater mitochondrial mass. In addition to calculating the ORR, factoring the side scatter (SSC-A) parameter in the optical redox ratio could provide a more quantitative estimate (aORR) of cellular metabolic capacity by accounting for differences in mitochondrial mass between cells, considering that mitochondria organelles are effective light scatters (Fig. 2C-E) ^16,17^.

**Figure 2:**
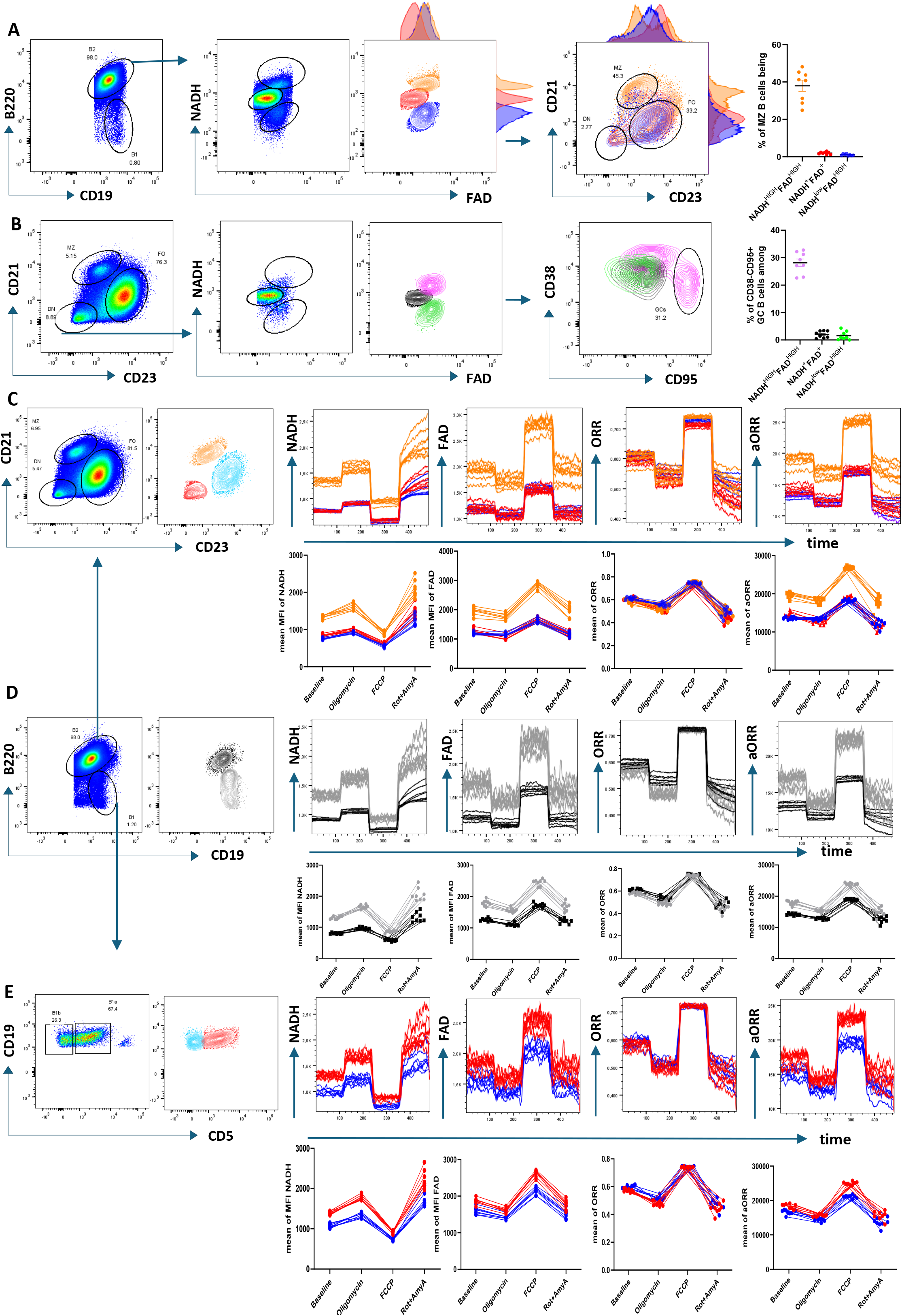
Measurement of NADH and FAD autofluorescence in the ultraviolet and blue region, respectively, with the FACSymphony A5 SE, in splenic B cell subpopulations from 18–25 weeks old male wild-type C57BL/6 mice (n=8). (A) NADH/FAD ratio analysis of B220^+^CD19^+^ B cells identified three metabolically distinct populations: a hypermetabolic NADH^HIGH^FAD^HIGH^ subset, a normometabolic NADH^+^FAD^+^ subset, and a NADH^LOW^FAD^HIGH^ subset. Backgating onto CD21 versus CD23 expression revealed that the hypermetabolic population predominantly localizes to the marginal zone (MZ) B cell compartment (p < 0.01). (B) Within the CD21^−^CD23^−^ double-negative (DN) B cell fraction, further stratification based on NADH and FAD identified three additional metabolic subsets, among which the NADH^HIGH^FAD^HIGH^ population corresponded significantly to activated CD38^HIGH^CD95^+^ B cells and newly formed CD38^−^CD95^+^ germinal center (GC) B cells (p < 0.01). See Supplementary Figure 1 for the gating strategy. (C–E) Flow cytometry-based real-time kinetics measurement of NADH and FAD, during application of the Mitostress test across (C) marginal zone (MZ), follicular (FO), and double negative (DN) B cells, (D) B2, and B1 B cells, and (E) CD5^+^ B1a and CD5^−^ B1b subsets, revealed pronounced intra-B cell metabolic heterogeneity. The data are presented as the means□± □SEMs and are representative of two independent experiments (n = 7-8). Statistical analysis: one-way ANOVA was conducted, when appropriate.

Advances in high-resolution spectral flow cytometry now allow integration of autofluorescence-based measurements with high-dimensional immunophenotyping, facilitating the analysis of metabolic heterogeneity across immune cell subsets *ex vivo*. Here, we present an optimized approach for multi-parameter autofluorescence-based metabolic measurements to interrogate metabolic heterogeneity across immune cell populations, with flow cytometry. Using the FACS Symphony A5 SE, we quantified NADH and FAD autofluorescence in major splenic B subsets and identified metabolically distinct immune cell subsets and activation states. We first applied this approach to major splenic B cell subsets (Figure 2 and Suppl. Figure 1) and calculated the *optical redox ratio* (ORR), defined as FAD / [NADH + FAD])^11,12^. In line with previous reports^18–21^, activated and highly proliferative germinal center B cells, and marginal zone B cells exhibited distinct autofluorescence signatures indicative of elevated metabolic activity compared with follicular B cells (Fig. 2 A-C). Furthermore, B1 cells display a more metabolically active phenotype than conventional B2, driven by CD5^+^ B1a cells, which exhibited higher autofluorescence-derived metabolic parameters than B1b cells (Fig. 2D, E). Comparison of the normalized ORR with the SSC-A-adjusted absolute ORR indicated that the hypermetabolic phenotype observed in innate-like MZ and B1 cells is primarily attributable to increased mitochondrial mass rather than enhanced activity per mitochondrion (Fig. 2 C-E).

## Data availability statement

All data generated and analysed during this research are available upon reasonable request.

## Permission to reproduce material from other sources

We acquired permission to reproduce Figure 1A from https://www.becker-hickl.com/applications/metabolic-imaging/, upon written request to Becker-Hickl

## Author contribution

ES conceptualized the project, performed experiments, analysed data, prepared figures, and wrote the original manuscript. NH supervised and reviewed the project, reviewed and edited the manuscript, and acquired and provided funding.

## Text editing

To improve clarity and readability, selected sections of the manuscript were refined using ChatGPT. The tool was used solely to assist with language editing after the initial draft had been prepared by the authors and did not contribute to the study design, scientific content, data analysis, interpretation of results, or conclusions. All AI-assisted revisions were critically reviewed and approved by the authors before submission.

## Acknowledgments and funding

We would like to thank Prof. Dr. med. Axel Methner for providing scientific input and advice. Also, we would like to thank CFFC manager, Kristian Schuetze, and the FACS Core Facility CFFC PKZI FZI of the University Medical Center of the Johannes Gutenberg-University Mainz for providing support and instrumentation of FACSymphony A5 SE funded by the Deutsche Forschungsgemeinschaft (DFG, German Research Foundation) - INST 371/45-1 FUGB.

N.H. was supported by the DFG grant HO4440/1-1, DFG grant HO4440/1-2, and Research Center Immunology (FZI) and further supported by the German Research Foundation (Project Number 318346496, SFB1292/2 TP20), and TRR355 TPA05.

---

**Supplementary Figure 1:**
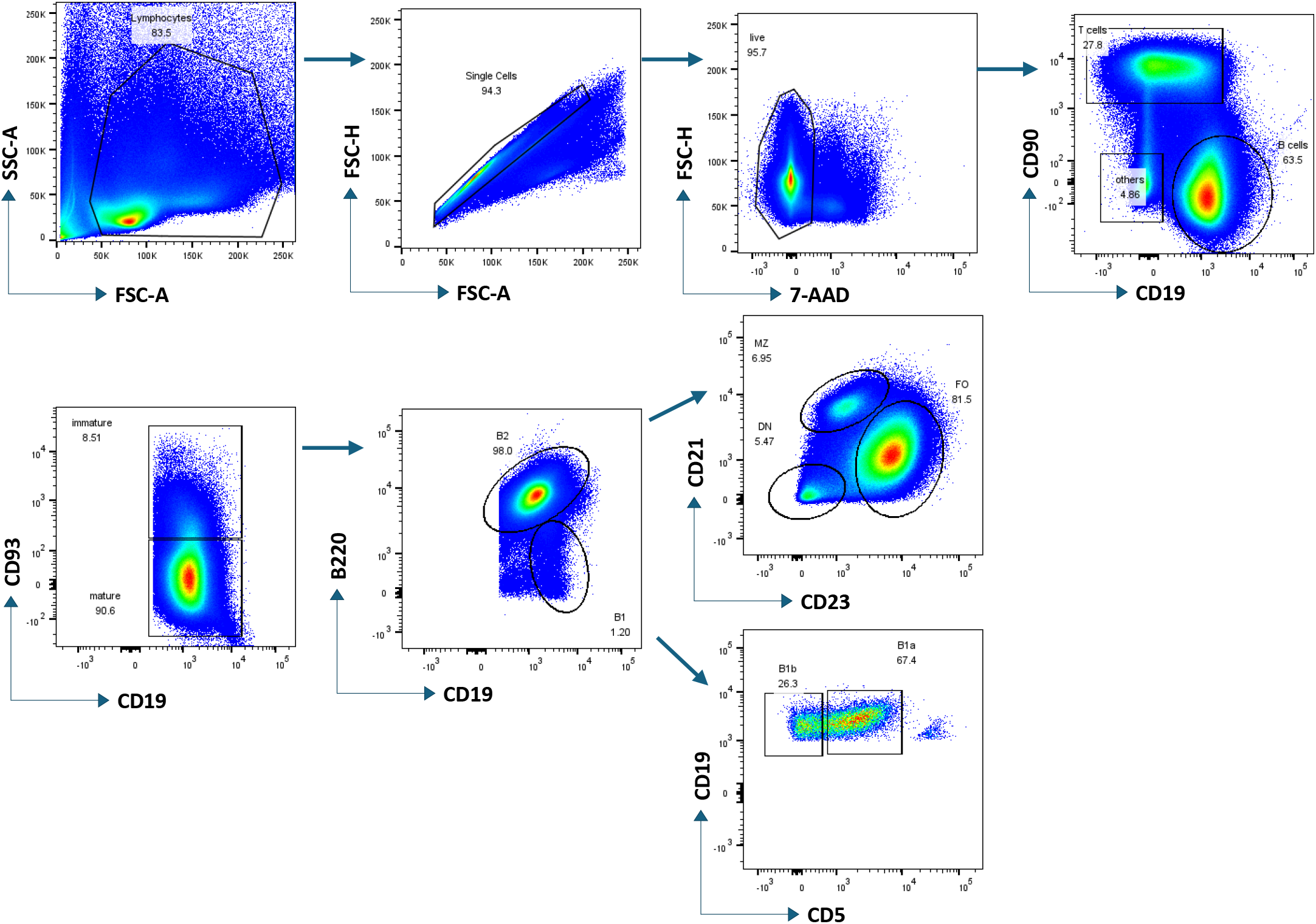
Gating strategy used for B cell metabolic profiling from splenic B cells

