## Supplementary material for "A streamlined spectral cytometry method for FAD and NADH autofluorescence analysis in immunometabolic studies": Materials and Methods

### Material and Methods

***Mice***. All the mice used were wild type on the C57BL/6J background, and they were bred in our facility (Translational Animal Research Center of the University Medical Center Mainz) under specific pathogen-free (SPF) conditions and kept at the experimental room facility. All animal experiments were performed in accordance with the institutional regulations of the Central Animal Facility.

***Preparation of single cell suspension for flow cytometry.*** To prepare a single cell suspension, spleens of C57BL/6 mice were smashed through a 40μm filter in FACS buffer (PBS buffer supplemented with 2% fetal calf serum). Cells were centrifuged at 300g for 7 minutes and resuspended in 1mL od ACK lysis buffer to lyse erythrocytes for approximately 3 minutes. Lysis reaction was terminated by adding 10mL of FACS buffer. After centrifugation, cells were resuspended in FACS buffer.

***Surface staining for analysis of B cell subpopulations***

Approximately 3-5x$10^6$ spleen cells were seeded on a 96-well U plate. Following centrifugation at 300g for 6 minutes, The Fc receptors were locked with anti-mouse CD16/32 (BioLegend) for 15 min at 4 °C. Following washing with FACS buffer, cells were stained with an antibody cocktail in FACS buffer (Table S1) for 20-30 minutes at 4oC in the dark. Cells were then washed twice in FCS-free DMEM medium (Thermo fisher), centrifuged and resuspended in glucose-containing FCS-free DMEM medium at a final volume of 500 µL in the presence of viability dye 7-AAD (BD Bioscience), and were left to equilibrate at room temperature to prevent temperature-induced metabolic shifts.

**Sample acquisition and measurement**

Samples were acquired at baseline, followed by sequential additions of oligomycin (20uM), FCCP (10uM), and rotenone/antimycin A (2uM) at approximately 1-minute intervals and a short, gentle vortex. Samples were analysed with a FACS Symphony A5 spectral unmixer (BD Bioscience). FCS files were analysed using the FlowJo v10.10 software. The antibody panels were designed as such to minimize spill-over into the channels used to measure NADH and FAD autofluorescence excited by the 355-nm UV laser and 488-nm Blue laser respectively. To augment autofluorescence intensity, detector voltages were incrementally increased to ensure that signals remained within the positive range. To circumvent spill-over issues of NADH to the FAD signal from UV excitation, FAD was excited with a 488-nm blue laser, which does not excite NADH. The ORR was calculated as [FAD/(NADH+FAD)], To further estimate absolute metabolic activity per cell, the side scatter was factored into ORR, resulting in an absolute ORR (aORR = ORR x SSC-A), based on the conjecture that mitochondrial mass correlates with cell size.

***Antibodies used for flow cytometry.*** The antibodies used for flow cytometry are listed in Table 1 and used as described in the Materials and Methods part of the main manuscript for the staining of viable cell suspensions.

Table S1: B cell panel for metabolic profiling

| **No** | **Fluorochrome conjugate** | **Markers** | **Clone** | **Manufacturer (Catalogue number)** |
|---|---|---|---|---|
| **1** | **BUV737** | **CD5** | 53-7.3 | BD Biosciences (612809) |
| **2** | **APC** | **CD93** | AA4.1 | eBioscience (17-5892) |
| **3** | **APC-Cy7** | **CD21** | **7E9** | BioLegend (123418) |
| **4** | **PE-Cy7** | **CD23** | B3B4 | eBioscience ( 25-0232) |
| **5** | **BV650** | **CD95** | Jo2 | BD Bioscience (740507) |
| **6** | **RB780** | **CD90** | 53-2.1 | BD Bioscience (755826) |
| **7** | **APC-R700** | **CD19** | 1D3 | BD Biosciences (565473) |
| **9** | **BV786** | **CD45R/B220** | RA3-6B2 | Biolegend (103246) |
| **10** | **RY610** | **CD38** | 90/CD38 | BD Biosciences (759659) |
| **11** | **-----------** | **7AAD viability dye** | **---------** | BD Biosciences (559925) |
